# CAR Macrophages for SARS-CoV-2 Immunotherapy

**DOI:** 10.1101/2020.07.26.222208

**Authors:** Wenyan Fu, Changhai Lei, Kewen Qian, Zetong Ma, Tian Li, Fangxin Lin, Wei Zhang, Jian Zhao, Shi Hu

## Abstract

Targeted therapeutics for the treatment of coronavirus disease 2019 (COVID-19), especially severe cases, are currently lacking. As macrophages have unique effector functions as a first-line defense against invading pathogens, we genetically armed human macrophages with chimeric antigen receptors (CARs) to reprogram their phagocytic activity against SARS-CoV-2. After investigation of CAR constructs with different intracellular receptor domains, we found that although cytosolic domains from MERTK (CAR_MERTK_) did not trigger antigen-specific cellular phagocytosis or killing effects, unlike those from MEGF10, FcRγ and CD3ζ did, these CARs all mediated similar SARS-CoV-2 clearance in vitro. Notably, we showed that CAR_MERTK_ macrophages reduced the virion load without upregulation of proinflammatory cytokine expression. These results suggest that CAR_MERTK_ drives an ‘immunologically silent’ scavenger effect in macrophages and pave the way for further investigation of CARs for the treatment of individuals with COVID-19, particularly those with severe cases at a high risk of hyperinflammation.

## Introduction

The coronavirus disease 2019 (COVID-19) pandemic has caused a sudden significant increase in hospitalizations for pneumonia with multiorgan disease and has led to more than 300,000 deaths worldwide. COVID-19 is caused by the novel severe acute respiratory syndrome coronavirus 2 (SARS-CoV-2), a novel enveloped RNA betacoronavirus. SARS-CoV-2 infection may be asymptomatic or cause a wide spectrum of symptoms, ranging from mild symptoms of upper respiratory tract infection to life-threatening sepsis.^1^ Manifestations of COVID-19 include asymptomatic carriers and fulminant disease characterized by sepsis and acute respiratory failure. Approximately 5% of patients with COVID-19, including 20% of those hospitalized, experience severe symptoms necessitating intensive care. More than 75% of patients hospitalized with COVID-19 require supplemental oxygen.^1,2^ The case-fatality rate for COVID-19 varies markedly by age, ranging from 0.3 deaths per 1000 patients among patients aged 5 to 17 years to 304.9 deaths per 1000 patients among patients aged 85 years or older. Among patients hospitalized in the intensive care unit, the case fatality can reach 40%.^1^

There is currently no human vaccine available for SARS-CoV-2, but approximately 120 candidates are under development. In the development of an effective vaccine, a number of challenges must be overcome, such as technical barriers, the feasibility of large-scale production and regulation, legal barriers, the potential duration of immunity and thus the number of vaccine doses needed to confer immunity, and the antibody-dependent enhancement effect. Moreover, there is another complicated area to consider: drug development for COVID-19, especially treatments for patients with severe or late-stage disease. Dexamethasone therapy was reported to reduce 28-day mortality in patients requiring supplemental oxygen compared with usual care (21.6% vs 24.6%; age-adjusted rate ratio, 0.83 [95% CI, 0.74-0.92])^3^, and remdesivir was reported to improve the time to recovery (hospital discharge or no supplemental oxygen required) from 15 to 11 days.^4^ In a randomized trial of 103 patients with COVID-19, convalescent plasma did not shorten the time to recovery^5^. Ongoing trials are testing antiviral therapies, immune modulators, and anticoagulants; however, there is no specific antiviral treatment recommended for COVID-19.

Chimeric antigen receptors (CARs) are synthetic receptors that redirect T cell activity towards specific targets^6^. A CAR construct includes antigen-recognition domains in the form of a single-chain variable fragment (scFv) or a binding receptor/ligand in the extracellular domains, a transmembrane domain providing the scaffold and signaling transduction, and intracellular domains from the T cell receptor (TCR) and costimulatory molecules that trigger T cell activation^7^. Based on the longstanding interest in harnessing macrophages to combat tumor growth8,9, human macrophages engineered with CARs have been developed and characterized for their antitumor potential. Macrophages, critical effectors of the innate immune system, are responsible for sensing and responding to microbial threats and promoting tissue repair. We therefore hypothesize that CAR macrophages can be used to combat SARS-CoV-2. However, the hyperinflammatory macrophage response, which has been found to be damaging to the host, particularly in severe infections, including SARS-CoV-2, and cytokine release syndrome (CRS), which is also the most significant complication associated with CAR-T cell therapy, raise questions regarding the safety of using CAR macrophages for virus clearance.

In this report, we developed a series of chimeric antigen receptors based on recognition of the S protein and tested their ability to induce phagocytosis of SARS-CoV-2 virions. Interestingly, we reported that one CAR with the intracellular domain of MERTK, which belongs to the TAM receptor family, did not show a notable killing effect in antigen-expressing cell-based models compared with other CARs but did demonstrate antigen-specific clearance of SARS-CoV-2 virions in vitro without the secretion of proinflammatory cytokines.

## Results

To program engulfment based on recognition of the SARS-CoV-2 spike protein, we used a CAR design for the synthetic receptor strategy in our study. The synthetic receptors were constructed to contain an scFv derived from an antibody recognizing the virus spike protein, CR3022, which has been reported to bind with the receptor-binding domain of the SARS-CoV-2 S glycoprotein with high affinity, and the CD8 transmembrane domain present in the αCD19 CAR for T cells^9^. For the cytoplasmic domains, we used the common γ subunit of Fc receptors (CARγ), MEGF10 (CAR_MEGF10_), MERTK (CAR_MERTK_) and CD3ζ (CARζ) in our study (Fig. 1a). These cytoplasmic domains are capable of promoting phagocytosis by macrophages.

**Figure 1.**
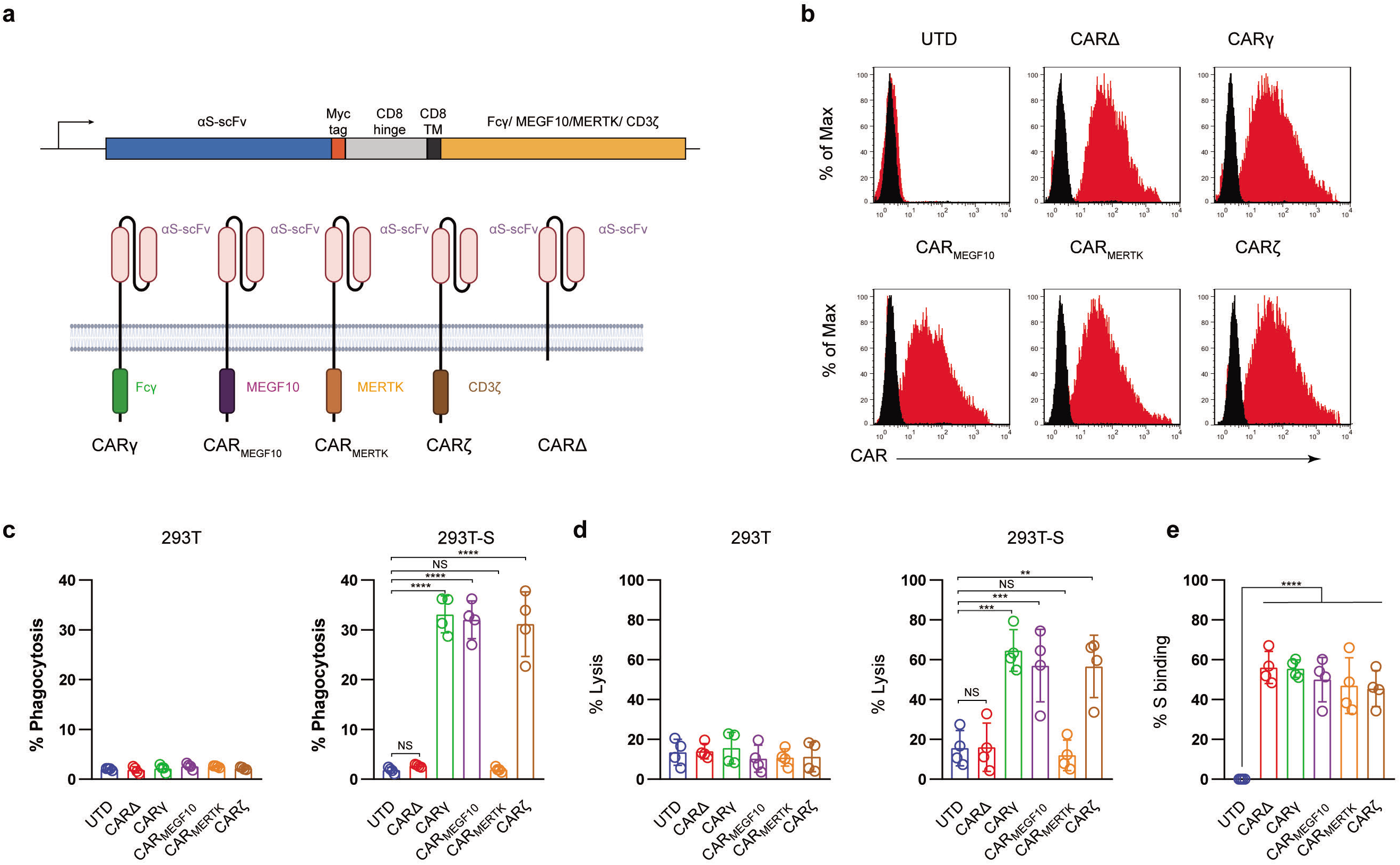
Generation and characterization of CAR macrophages. **a**, Vector maps of tested CAR designs and schematics showing the structures of CARs used in the study. Figure created with BioRender. **b**, Membrane-bound CAR expression. Forty-eight hours after retroviral transduction, the expression of synthetic receptors on THP-1 cells was detected by staining with an anti-MYC antibody, followed by flow cytometry analysis. Untransduced THP-1 cells were used as a negative control. The histograms shown in black correspond to the isotype controls, whereas the red histograms indicate positive fluorescence. **c**, FACS-based phagocytosis of 293T cells or 293T-S target cells by UTD or different CAR macrophages. Statistical significance was calculated with one-way ANOVA with multiple comparisons, and data represent n = 3 technical replicates (representative of at least three individual experiments). **d**, Killing of 293T or 293T-S cells by UTD or anti-S CAR macrophages at 24 h assessed with a luciferase-based assay. **e.** Flow cytometry analyses of CAR macrophages stained with a biotinylated S protein followed by streptavidin-FITC. The histograms shown in black correspond to the use of isotype controls with streptavidin-FITC, whereas the red histograms indicate positive fluorescence. The results shown represent three (b) independent experiments. Data are the shown as the mean ± s.d. of four independent biological replicates (c, d, e). P values were derived by one-way ANOVA followed by Tukey’s posttest (c, d, e). *p<0.05, **p<0.01, ***p<0.001, ****p<0.0001.

Next, we used lentiviral vector technology to express the fusion constructs in human macrophage THP-1 cells using clinically validated techniques^10^. The cDNA sequences containing the various fusion constructs were cloned into a third-generation lentiviral vector in which the CMV promoter was replaced with the EF-1α promoter^11^. An extracellular MYC epitope was cloned into the receptors to permit detection by flow cytometry. Lentiviral vector supernatants transduced THP-1 cells with high efficiency (Fig. 1b). The phagocytic potential of human macrophage THP-1 cell lines expressing different CAR receptors or a truncated CAR receptor (CARΔ) lacking the intracellular domain was measured with a cell-based assay. Consistent with previous reports8,9, CAR macrophages and control untransduced (UTD) macrophages did not show notable phagocytosis of 293 cells; however, CAR_MEGF10_, CARγ and CARζ cells but not CAR_MERTK_, CARΔ, or UTD macrophages phagocytosed Spike-bearing 293 cells in an S-specific manner (Fig. 1c). CAR-mediated macrophage phagocytosis was further confirmed by a luciferase-based killing assay, and our data showed that CAR_MEGF10_, CARγ and CARζ cells eradicated S protein-expressing 293T cells in an antigen-specific manner (Fig. 1d). Interestingly, CAR_MERTK_ and UTD macrophages showed-no difference in killing effect. Our data further showed that all synthetic receptors had the ability to bind the S protein (Fig. 1e); therefore, the differences in phagocytosis and the lytic effect were not due to the affinity for the S protein.

Although there is currently no evidence that SARS-CoV-2 can infect THP-1 cells with or without IgGs^12^, THP-1 cells have been shown to support antibody-mediated enhancement of SARS-CoV infection in previous studies^13^. We therefore sought to determine whether synthetic receptors facilitate the entry of SARS-COV-2 into macrophages as host cells, as the extracellular domain of the CAR constructs has the capacity to directly bind to the S protein. Replication-defective VSV particles bearing coronavirus S proteins faithfully reflect key aspects of host cell entry by coronaviruses, including SARS-CoV-2^14,15^. We therefore employed VSV pseudotypes bearing SARS-2-S to study the cell entry of SARS-CoV-2. Our data showed that Vero E6 cells were susceptible to entry driven by SARS-S (Fig. 2a); however, no evidence of infection was detected in THP-1 cells with or without synthetic receptors.

**Figure 2.**
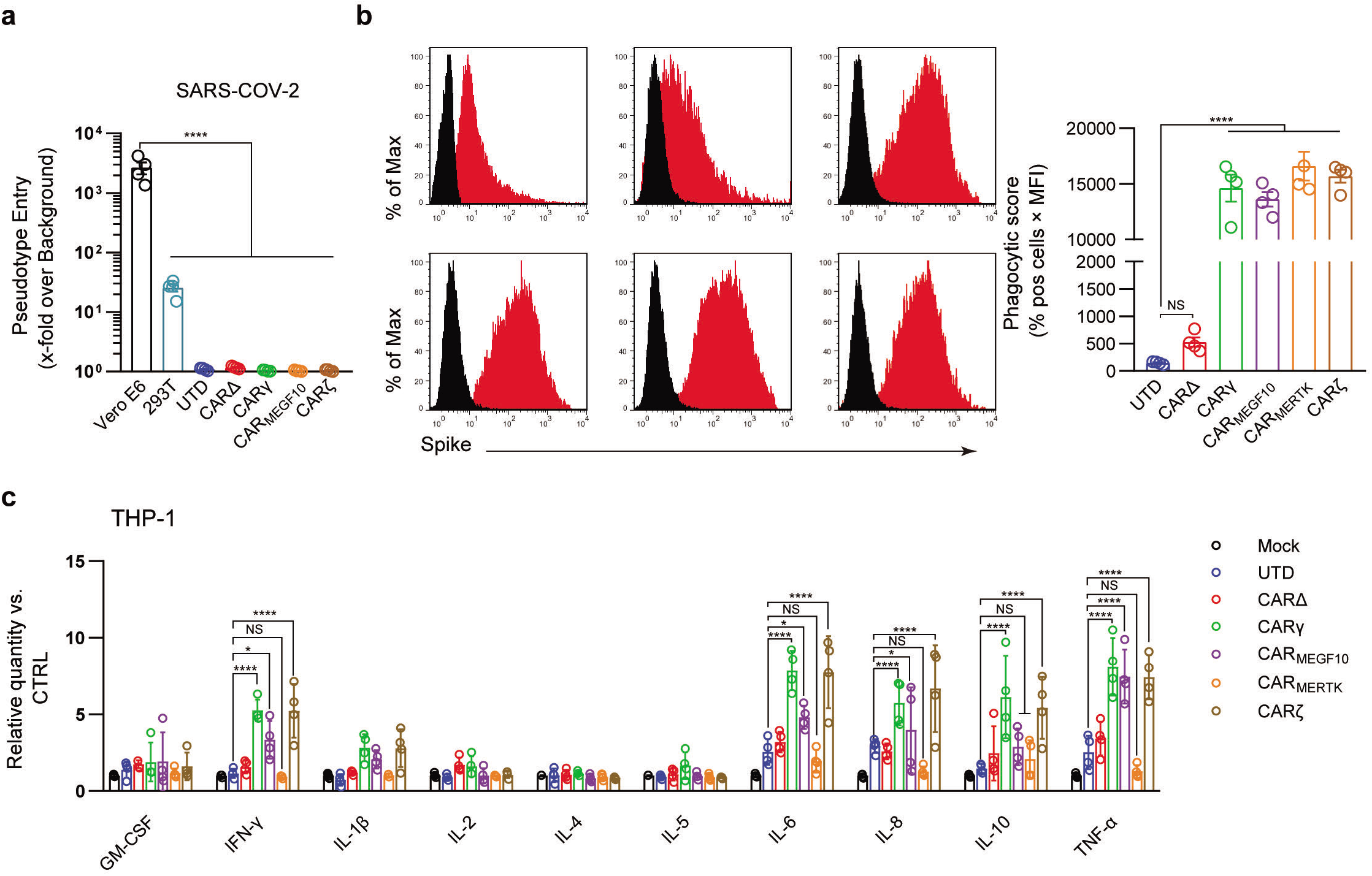
CARs mediate phagocytosis of SARS-CoV-2 virions. **a,** Different cell lines were inoculated with a SARS-CoV-2 pseudotyped virus. At 16 h postinoculation, pseudotyped virus entry was analyzed by determining the luciferase activity in cell lysates. Signals obtained for particles bearing no envelope protein were used for normalization. The average of three independent experiments is shown. Error bars indicate the SEM. **b,** The uptake of pseudotyped virions by UTD and CAR macrophages was analyzed by flow cytometry. Different cell lines were stained with an anti-S primary Ab. The histograms shown in black correspond to the isotype controls, whereas the red histograms indicate positive fluorescence. Data are reported as the phagocytic score (% positive cells x MFI, right panel). **c**, Cell lines were infected with the SARS-CoV-2 pseudotyped virus or mock infected. Cytokine levels in the supernatants were determined by a multiplex bead array. The relative level was calculated as the ratio of the infected cells to the mock-infected THP-1 cells. Data are shown as the mean ± s.d. (a–c) of four independent biological replicates. P values were derived by one-way ANOVA followed by Tukey’s posttest (a–b) or two-way ANOVA followed by the Bonferroni posttest (c); *p<0.05, **p<0.01, ***p<0.001, ****p<0.0001.

Antibody-mediated phagocytosis and internalization of virions are important mechanisms of antiviral activity performed by macrophages against pathogens; however, using the phagocytosis assay developed for SARS-CoV-2, we observed low levels of phagocytic activity when UTD cells directly contacted virions. Phagocytic activity was not significantly increased when CARΔ cells rather than UTD macrophages were the phagocytes in the assay, suggesting that the extracellular domain of the CAR alone is not sufficient to induce strong virion internalization. CARγ, CAR_MEGF10_, and CARζ mediated similar significantly stronger levels of SARS-CoV-2 phagocytosis by THP-1 cells than CARΔ (Fig 2b). Unexpectedly, we also observed strong internalization of virions in CAR_MERTK_ cells, which did not show specific phagocytic or lytic effects on S protein-expressing 293T cells. Since all the CARs exhibited the ability to induce phagocytosis of SARS-CoV-2 virions while there was no evidence of infection, these experiments strongly suggest the clearance of SARS-CoV-2 virions of CAR macrophages.

Because the systemic cytokine profiles observed in patients with severe COVID-19 show similarities to those observed in patients with macrophage activation syndrome, culture supernatants from THP-1 cells with different CARs treated with virions were further analyzed in a multiplex cytokine assay (Fig. 2c). Following SARS-CoV-2 treatment of THP-1 cells, we observed slightly increased secretion of the cytokines IL-6, IL-8 and TNF-α, but no discernable patterns could be confidently drawn for GM-CSF, IL-1β, IL-2, IL-4, IL-5, IL-8, IL-10, and IFN-γ. CARΔ cells showed a cytokine profile similar to that of UTD macrophages. Notably, we observed not only strongly increased induction of IL-6, IL-8 and TNF-α but also induction of IFN-γ and IL-10 in SARS-CoV-2-treated CARγ and CARζ cells. However, for CAR_MERTK_ cells, we did not observe significant changes in cytokines.

We further used a transwell-based coculture model to evaluate the protective role of CAR macrophages in SARS-CoV-2 infection (Fig. 3a). All the CAR-expressing macrophages potently inhibited Vero E6 cell infection with the SARS-CoV-2-S pseudotyped virus. Interestingly, CARΔ cells showed no protective effect in the infection assay, although they had a similar capacity to bind to the S protein, suggesting that the intracellular signaling domain is necessary for virion clearance by CAR macrophages (Fig. 3b).

**Figure 3.**
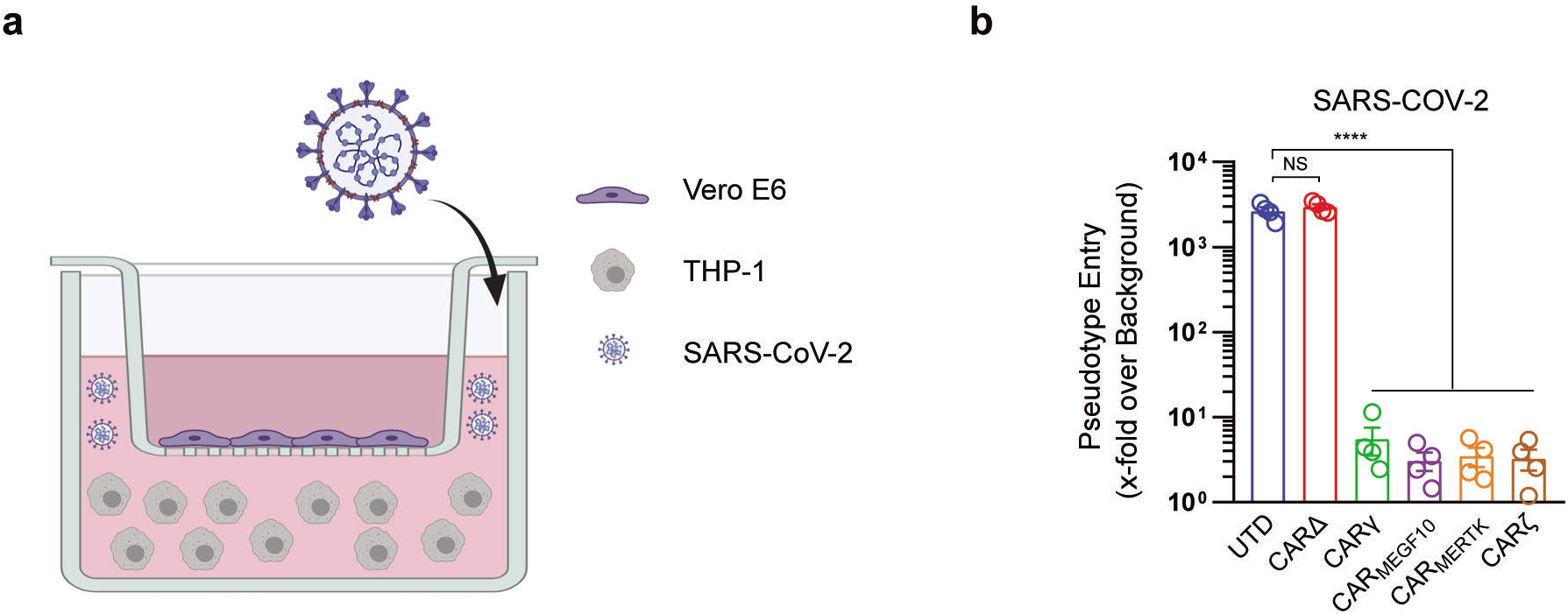
CARs mediate protection against SARS-CoV-2 infection. **a**, The schematic shows the transwell coculture model. Figure created with BioRender. **b,** Different cell cocultures were inoculated with the SARS-CoV-2 pseudotyped virus in the culture plate. At 16 h postinoculation, pseudotyped virus entry was analyzed by determining the luciferase activity in cell lysates. Signals obtained for particles bearing no envelope protein were used for normalization. Data are presented as the mean ± s.d. (a–c) of four independent biological replicates. P values were derived by one-way ANOVA followed by Tukey’s posttest. *p<0.05, **p<0.01, ***p<0.001, ****p<0.0001.

## Discussion

Macrophages, which protect against infections and scavenge the body’s worn-out or abnormal cells, are known for their phagocytic activity, antigen presentation capability, and flexible phenotype. The innate immune response of the pulmonary parenchyma, which is characterized by the differentiation of bone marrow-derived monocytes into macrophages, serves as a first-line defense against invading pathogens in the lungs^16^. In general, monocytes/macrophages are able to remarkably limit viral replication. The monocyte-enhanced proinflammatory signaling molecule levels and antiviral responses provoked during viral infection have been shown for influenza, herpes, and Zika viruses^17^. Moreover, it has recently been suggested that some COVID-19 patients have enhanced proinflammatory macrophage activity, which leads to accelerated production of inflammatory cytokines and chemokines and has mostly been observed in subjects with a poor prognosis^18^.

To our knowledge, no synthetic cell-based immunotherapy has been investigated for COVID-19. CAR-expressing T cells have been demonstrated to be a very effective approach to treat B-cell cancer patients. Harnessing the power of engineered macrophages for the development of novel treatments for solid tumors is of great interest because CAR-T cell therapy is often hampered by the inability of T cells to penetrate solid tumors and the inhibitory tumor microenvironment^19^. Consistent with a previous report^8^, CAR receptors with cytosolic immunoreceptor tyrosine-based activation motifs (ITAMs) were capable of triggering specific engulfment and killing of antigen-expressing cells by macrophages. These CAR macrophages also showed strong phagocytosis of SARS-COV-2 virions in our data; however, this effect was accompanied by increased secretion of the proinflammatory cytokines IFN-γ, IL-6, and IL-8. In CAR-T cell therapy, engineered T cell expansion is usually accompanied by high-grade CRS with elevated circulating levels of interferon (IFN)-γ, granulocyte-colony stimulating factor (G-CSF), IL-6, IL-8 and IL-10. Recent reports have demonstrated that host-derived monocyte/macrophage and CAR-T cell interactions play an important role in CRS pathophysiology^20^. This is of interest because increased serum levels of similar inflammatory cytokines^21–23^ have been associated with COVID-19 severity and death. Interestingly, the secretion of IL-6, IL-8, TNF-α, IFN-γ and IL-10 was significantly elevated in CARγ and CARζ cells treated with SARS-CoV-2 virions, suggesting that these CAR macrophages may not be suitable for application in severe patients or patients with late-stage COVID-19.

Previous studies have shown that human immune cells, such as THP-1 cell lines, are susceptible to SARS-CoV infection^24^. We did not observe any evidence that our SARS-CoV-2 pseudotyped virus infected THP-1 cells. Moreover, the uptake of virions by THP-1 cells was very low, even with a truncated CAR with the ability to bind to the S protein, suggesting that THP-1 cells did not innately engulf the virions. Notably, CAR_MERTK_, which was regarded as an unsuccessful receptor in a previous report^8^ and showed no cellular killing effect on target cells when expressed in THP-1 cells in our assay, demonstrated a virion clearance capacity similar to that of CARγ and CARζ. Our data further support that CAR_MERTK_ mediates ‘immunologically silent’ virion removal, which does not elicit a proinflammatory response.

MER tyrosine kinase (MERTK), together with TRYO3 and AXL, belongs to the TAM family of receptor tyrosine kinases (RTKs). These receptors can be activated by a complex ligand consisting of phosphatidylserine (PtdSer) linked to the RTK by a vitamin K-dependent protein ligand, Gas6, or Protein S^25^, playing a crucial role in innate immune cells. Gas6 has the capacity to bind all three receptors, while Protein S is a specific ligand of MERTK and TYRO3^26^. Apoptotic cells, exosomes, and cell debris are the main sources of the PtdSer component. In some cases, the PtdSer component is also provided by patches of exposed PtdSer on living cells (including T cells)^25^. The activation of members of the TAM family of receptors generally induces an anti-inflammatory, homeostatic response in innate immune cells, diminishing excessive inflammation and autoimmune responses elicited by the ingestion of “self’^25^. However, previous studies also proposed that enveloped viruses may hijack TAM receptors to facilitate attachment and infection via a PtdSer-dependent process termed “apoptotic mimicry” and act as potent TAM agonists, in turn inhibiting the type I IFN response in target cells^27^. In our study, THP-1 cells expressing the synthetic receptor with the MERTK cytoplasmic domain were relatively resistant to virus infection but induced notable virion clearance. It should be noted that our study used very simple infection models; therefore, the assays lack numerous physiological and pathological factors, such as IgG or complement-mediated immune complexes, that may interfere with the behavior of engineered cells. Of cause, cells expressing synthetic receptors can be further engineered and developed to achieve precise control.

In summary, our data reveal that the CAR-based synthetic approach is applicable for COVID-19 treatment. In addition to direct virion clearance by CAR macrophages, we found evidence that MERTK-based CAR receptors did not induce further upregulation of proinflammatory cytokine levels, thereby raising the possibility that CAR macrophages may be useful as potent therapeutics in severe COVID-19.

## Methods

### Cell lines

All cell lines were purchased from the American Type Culture Collection (ATCC; Manassas, VA). The identities of the cell lines were verified by STR analysis, and the cell lines were confirmed to be mycoplasma free. 293 and Vero cells were maintained in DMEM supplemented with 10% fetal bovine serum, and THP-1 cells were maintained in RPMI medium supplemented with 10% fetal bovine serum. Cell culture media and supplements were obtained from Life Technologies, Inc.

### Vector construction

The sequence encoding the scFv generated from CR3022 was chemically synthesized. As shown in Fig. 1a, synthetic receptors contained the human CD8α signal peptide followed by the scFv linked in-frame to the hinge domain of the CD8α molecule, transmembrane region of the human CD8 molecule, and intracellular signaling domains of the FCER1G, MEGF10, MERTK or CD3ζ molecules. The fragments were subcloned into the pELNS vector^28^. High-titer replication-defective lentiviruses were produced and concentrated^28^. Lentiviral infection was used to stably express CAR constructs in THP-1 cells.

### FACS-based phagocytosis assay

UTD or CAR-expressing THP-1 cells were cocultured with GFP^+^ 293T cells or GFP^+^ 293T-S (S^+^) target cells for 4 h at 37 °C. The effector-to-target (E:T) ratio was 1:1, and 1 × 10^5^ cells were used as both effector cells and target cells. After coculturing, the cells were harvested and stained with an anti-CD11b APC-Cy7-conjugated antibody (M1/70, BioLegend) and analyzed by FACS using a FACSCalibur flow cytometer (BD Biosciences). The percentage of phagocytosis was calculated based on the percent of GFP^+^ events within the CD11b^+^ population. Data are represented as the mean ± standard error of quadruplicate wells.

### Flow cytometry

Cell-surface staining was performed for 45 min at 4 °C and was analyzed using a FACSCalibur flow cytometer (BD Biosciences). A minimum of 1 × 10^4^ events per sample were examined.

### In vitro cytotoxicity assay

293T and 293T-S cells were used as targets in luciferase-based killing assays including control (UTD) or CAR macrophages. The effector-to-target (E:T) ratio was 10:1 for all the groups. Bioluminescence was measured using a Bio-Tek Synergy H1 microplate reader. The percent specific lysis was calculated on the basis of the experimental luciferase signal (total flux) relative to the signal of the target alone, using the following formula: %Specific Lysis = [(Sample signal − Target alone signal)] / [Background signal − Target alone signal)] × 100.

### SARS-CoV-2 pseudovirus and cell infection experiments

The SARS-CoV-2 pseudovirus was constructed based on the spike genes of the strain Wuhan-Hu-1 (GenBank: MN908947) using published methods^29^. The SARS-CoV-2 spike gene was chemically synthesized and cloned into a eukaryotic expression plasmid. 293T cells were first transfected with the S expression vector and then infected with a VSV pseudotyped virus (G*ΔG-VSV), in which the VSV-G gene was substituted with luciferase expression cassettes. The culture supernatants were harvested and filtered at 24 h postinfection. The SARS-CoV-2 pseudovirus could not be neutralized with anti-VSV-G antibodies, and no G*ΔG-VSV was mixed with the SARS-CoV-2 pseudovirus stock. For cell-based infection assays, target cells were grown in plates until they reached 50%–75% confluency and then were inoculated with pseudotyped virus. The transduction efficiency was quantified at 16 h posttransduction by measuring firefly luciferase activity according to the manufacturer’s instructions (Promega).

### Phagocytosis assay

In all cases, SARS-CoV-2 S pseudotyped virions were pelleted (90 min at 14,000 rpm and 4 °C), and after removal of the supernatant, the pellets were resuspended in RPMI medium and incubated with phagocytes (THP-1 cells or CAR macrophages) at 37 °C for 1.5 h. After allowing time for phagocytosis, the cells were washed three times with PBS and incubated with Accutase (Innovative Cell Technologies) for 10 min at 37 °C, followed by a final wash in Accutase. Intracellular staining for the S protein was performed for 60 min on ice after using a fixation/permeabilization kit (eBioscience) and then analyzed using a FACSCalibur flow cytometer (BD Biosciences). The phagocytic score was determined by gating the samples on events representing cells and was calculated as follows: Percent S protein positive × median fluorescence intensity (MFI).

### Cytokine analysis

Cytokine analysis was performed on supernatants derived from cultures given the indicated treatments using a human cytokine 10-plex panel (Thermo Scientific) per the manufacturer’s instructions, with the panel results read on a Luminex Analyzer.

### Statistical analysis

Unless otherwise specified, Student’s t test was used to evaluate the significance of differences between two groups, and ANOVA was used to evaluate differences among three or more groups. Differences between samples were considered statistically significant when P < 0.05.

## Acknowledgements

This study was supported by the National Natural Science Foundation of China (grant nos. 82041012, 81773261, 31970882, 81903140 and 81602690); the Shanghai Rising-Star Program (grant no. 19QA1411400); and the Shanghai Sailing Program (19YF1438600).

## References

1 Wiersinga, W. J., Rhodes, A., Cheng, A. C., Peacock, S. J. & Prescott, H. C. Pathophysiology, Transmission, Diagnosis, and Treatment of Coronavirus Disease 2019 (COVID-19): A Review. JAMA, doi:10.1001/jama.2020.12839 (2020).

2 Garg, S. et al. Hospitalization Rates and Characteristics of Patients Hospitalized with Laboratory-Confirmed Coronavirus Disease 2019 - COVID-NET, 14 States, March 1-30, 2020. MMWR Morb Mortal Wkly Rep 69, 458–464, doi:10.15585/mmwr.mm6915e3 (2020).

3 Horby, P. et al. Effect of Dexamethasone in Hospitalized Patients with COVID-19: Preliminary Report. medRxiv, 2020.2006.2022.20137273, doi:10.1101/2020.06.22.20137273 (2020).

4 Beigel, J. H. et al. Remdesivir for the Treatment of Covid-19 - Preliminary Report. N Engl J Med, doi:10.1056/NEJMoa2007764 (2020).

5 Wang, W. et al. Detection of SARS-CoV-2 in Different Types of Clinical Specimens. JAMA, doi: 10.1001/jama.2020.3786 (2020).

6 June, C. H., O’Connor, R. S., Kawalekar, O. U., Ghassemi, S. & Milone, M. C. CAR T cell immunotherapy for human cancer. Science 359, 1361–1365, doi:10.1126/science.aar6711 (2018).

7 Fesnak, A. D., June, C. H. & Levine, B. L. Engineered T cells: the promise and challenges of cancer immunotherapy. Nature Reviews Cancer 16, 566–581 (2016).

8 Morrissey, M. A. et al. Chimeric antigen receptors that trigger phagocytosis. Elife 7, e36688 (2018).

9 Klichinsky, M. et al. Human chimeric antigen receptor macrophages for cancer immunotherapy. Nature Biotechnology, 1–7 (2020).

10 Levine, B. L. et al. Gene transfer in humans using a conditionally replicating lentiviral vector. Proceedings of the National Academy of Sciences 103, 17372–17377 (2006).

11 Fu, W. et al. CAR exosomes derived from effector CAR-T cells have potent antitumour effects and low toxicity. Nature communications 10, 1–12 (2019).

12 Banerjee, A. et al. Isolation, Sequence, Infectivity, and Replication Kinetics of Severe Acute Respiratory Syndrome Coronavirus 2. 26 (2020).

13 Jaume, M. et al. Anti-severe acute respiratory syndrome coronavirus spike antibodies trigger infection of human immune cells via a pH- and cysteine protease-independent FcγR pathway. Journal of virology 85, 10582–10597, doi:10.1128/JVI.00671-11 (2011).

14 Kleine-Weber, H. et al. Mutations in the spike protein of Middle East respiratory syndrome coronavirus transmitted in Korea increase resistance to antibody-mediated neutralization. 93 (2019).

15 Hoffmann, M. et al. SARS-CoV-2 cell entry depends on ACE2 and TMPRSS2 and is blocked by a clinically proven protease inhibitor. (2020).

16 Abassi, Z., Knaney, Y., Karram, T. & Heyman, S. N. The Lung Macrophage in SARS-CoV-2 Infection: A Friend or a Foe? Frontiers in Immunology 11, 1312 (2020).

17 Nikitina, E., Larionova, I., Choinzonov, E. & Kzhyshkowska, J. Monocytes and Macrophages as Viral Targets and Reservoirs. International journal of molecular sciences 19, doi:10.3390/ijms19092821 (2018).

18 Vaninov, N. In the eye of the COVID-19 cytokine storm. Nature reviews. Immunology 20, 277, doi:10.1038/s41577-020-0305-6 (2020).

19 Ritchie, D. et al. In vivo tracking of macrophage activated killer cells to sites of metastatic ovarian carcinoma. Cancer Immunol Immunother 56, 155–163 (2007).

20 Giavridis, T. et al. CAR T cell-induced cytokine release syndrome is mediated by macrophages and abated by IL-1 blockade. Nat Med 24, 731–738, doi:10.1038/s41591-018-0041-7 (2018).

21 Huang, C. et al. Clinical features of patients infected with 2019 novel coronavirus in Wuhan, China. The lancet 395, 497–506 (2020).

22 Chen, G. et al. Clinical and immunological features of severe and moderate coronavirus disease 2019. The Journal of clinical investigation 130 (2020).

23 Merad, M. & Martin, J. C. Pathological inflammation in patients with COVID-19: a key role for monocytes and macrophages. Nature Reviews Immunology, 1–8 (2020).

24 Yen, Y. T. et al. Modeling the early events of severe acute respiratory syndrome coronavirus infection in vitro. Journal of virology 80, 2684–2693, doi:10.1128/jvi.80.6.2684-2693.2006 (2006).

25 Graham, D. K., DeRyckere, D., Davies, K. D. & Earp, H. S. J. N. r. C. The TAM family: phosphatidylserine-sensing receptor tyrosine kinases gone awry in cancer. 14, 769–785 (2014).

26 Tsou, W.-I. et al. Receptor tyrosine kinases, TYRO3, AXL, and MER, demonstrate distinct patterns and complex regulation of ligand-induced activation. 289, 25750–25763 (2014).

27 Moller-Tank, S. & Maury, W. J. V. Phosphatidylserine receptors: enhancers of enveloped virus entry and infection. 468, 565–580 (2014).

28 Carpenito, C. et al. Control of large, established tumor xenografts with genetically retargeted human T cells containing CD28 and CD137 domains. 106, 3360–3365 (2009).

29 Nie, J. et al. Establishment and validation of a pseudovirus neutralization assay for SARS-CoV-2. 9, 680–686 (2020).

